# Seqwin: Ultrafast identification of signature sequences in microbial genomes

**DOI:** 10.1101/2025.11.07.687294

**Authors:** Michael X. Wang, Bryce Kille, Michael G. Nute, Siyi Zhou, Lauren B. Stadler, Todd J. Treangen

## Abstract

**Motivation:** Polymerase chain reaction (PCR) enables rapid, cost-effective diagnostics but requires prior identification of genomic regions that allow sensitive and specific detection of target microbial groups, herein referred to as microbial signature sequences. We introduce Seqwin, an open-source framework designed to automate microbial genome signature discovery. Tens of thousands of microbial genomes are now available for a single species, limiting the application of existing manual and automated approaches for identifying signatures. Modern approaches that are capable of leveraging all available microbial genomes will ensure sensitive and accurate DNA signature identification and enable robust pathogen detection for clinical, environmental, and public health applications.

**Results:** Seqwin builds weighted pan-genome minimizer graphs and uses a traversal algorithm to identify signature sequences that occur frequently in target genomes but remain rare in non-targets. Unlike earlier tools that depend on strict presence or absence of sequences, Seqwin accommodates natural sequence variation and scales to very large genome collections. When applied to genomes from C. difficile, M. tuberculosis, and S. enterica, Seqwin recovered more high-quality signatures than alternative methods with lower computational burden. Seqwin’s analysis of nearly 15,000 S. enterica genomes yielded over 200 candidate signatures in 5 minutes. Seqwin provides an open-source solution for the long-standing need for scalable microbial signature discovery and diagnostic assay design.

**Availability and Implementation:** Seqwin is freely available for academic use (https://github.com/treangenlab/Seqwin) and can be installed via Bioconda. Benchmarking datasets, outputs, and scripts are available on Zenodo https://doi.org/10.5281/zenodo.19176444.

**Supplementary Materials:** Provided as separate PDF and data files.

## Introduction

PCR-based infectious disease diagnostics have been the clinical standard for more than three decades, providing rapid and reliable detection of specific pathogens [1]. Clinical studies in the late 1980s, such as the detection of HIV-1 DNA in infants and adults [2], first demonstrated its utility in patient diagnosis. These early studies laid the foundation for PCR-based infectious disease diagnostics. Signature sequences are genomic regions that allow accurate identification and classification of microorganisms within specific taxonomic groups. DNA signature sequences were originally defined to be sequences both 100% conserved over all target genomes and unique compared to all non-target genomes [3]. However, over time the definition has been relaxed in signature detection tools and is now more commonly defined as a sequence that is highly sensitive (largely present in most target genomes), and highly specific (largely absent in or divergent to closely related non-target genomes).

Early computational methods for signature discovery mainly focused on microarray and TaqMan assay design, and they usually required signatures to be perfectly conserved in all target genomes [4, 5, 6, 7, 8]. Approaches such as YODA [4] and ProDesign [5] use exhaustive searches combined with various filtering criteria, while Insignia [6, 7] and CaSSiS [8] leverage maximal unique matches (MUMs) identified via a suffix tree data structure. While all of these methods were important advances in microbial signature discovery, they suffer from several limitations. First, tools developed for the design of microarray probes typically produce short signatures (≤50 bp) [4, 5], which are often insufficient for modern targeted sequencing assays. Second, many of these methods were developed before the NGS era, when genomic data was considerably limited. When Insignia pioneered open-source microbial signature discovery, only tens to hundreds of genomes were available for a given microbial target. Consequently, these tools were not designed to handle modern genomic databases at terabyte-to-petabyte scales. Furthermore, due to their reliance on exact matches across all target genomes, these methods are highly vulnerable to sequence variation, limiting their application in the context of increasingly diverse genomic datasets. On the other hand, non-exact, match-based approaches provide improved sensitivity but poor scalability. For example, SigSeekr [9] relies on BLAST-based [10] genome subtraction methods, but suffers from prohibitive runtime on large-scale datasets.

Recent years have seen a new wave of tools for genomic signature identification that emphasize scalability and flexibility. Many abandon all-versus-all sequence alignment in favor of ultrafast *k*-mer-based strategies or clever filtering techniques. Fur [11, 12] combines targeted genomic subtraction and stringent intersection methods to discover unique genomic regions. However, it struggles with highly divergent target genomes and/or similar background genomes, and may produce few or no candidate signatures. Methods such as Neptune [13], KmerGO [14], KEC [15] and Unikseq [16, 17] use alignment-free *k*-mer filtering and subsequently combine *k*-mers into longer signatures through sequence assembly or similar strategies. However, they all require substantial memory to store *k*-mers from all input genomes. NAUniSeq [18] is a recent *k*-mer-based method that automates the selection of target and non-target taxa, reporting identified *k*-mers directly as signatures without further extension. As a result, it typically requires a larger *k* (e.g., 100 bp), and all output signatures are fixed at that length. CURED [19] uses a similar *k*-mer filtering strategy and reports restriction sites on the output *k*-mers. While some of these tools do not explicitly use the term “signature”, they computationally address the same problem using target and non-target genome groups as input. Table 1 summarizes these tools, including their publication years and approaches. While advances in computational tools have provided more scalable solutions, each still grapples with trade-offs related to sensitivity and scalability on terabyte-sized datasets, motivating the development of solutions that can address this unmet need.

**Table 1.** Existing tools for identifying signature sequences in microbial genomes.

| Name | Year<sup>1</sup> | Approach |
| --- | --- | --- |
| Insignia [6, 7] | 2009 | Suffix tree |
| SigSeekr [9] | 2015 | BLAST |
| Neptune [13] | 2017 | $k$-mer assembly |
| KmerGO [14] | 2020 | $k$-mer assembly |
| KEC [15] | 2021 | $k$-mer assembly |
| Fur [11, 12] | 2024 | Sequence filtering pipeline |
| NAUniSeq [18] | 2024 | $k$-mer filtering |
| Unikseq [16, 17] | 2025 | $k$-mer assembly |
| CURED [19] | 2026 | $k$-mer filtering |
| Seqwin | 2026 | Minimizer graph |

<sup>1</sup> Year of the latest publication.

To address this gap, we developed Seqwin, a rapid and flexible algorithm for genomic signature discovery using pan-genome minimizer graphs. Unlike previous minimizer graph-based methods that require each minimizer to be present in all input genomes [20, 21], Seqwin allows the inclusion of minimizers from any of the input genomes and penalizes those absent in targets and/or present in non-targets. It then extracts connected low-penalty subgraphs and determines a representative sequence (signature) for each subgraph with a memory-efficient workflow. This novel minimizer graph approach increases the retention of original sequence information while keeping a low memory profile, and enables fast and flexible search of genomic signatures. In related work, mdBG constructs a minimizer-space de Bruijn graph for long-read assembly [22]; in contrast, Seqwin builds minimizer adjacency graphs (edges reflect consecutive minimizers along genomes) for comparative analyses.

Due to its efficient and flexible design, Seqwin performs well at identifying signatures from large collections of microbial genomes despite sequence variations, a critically important capability given the heterogeneity and rapid expansion of contemporary genomic databases. This allows Seqwin to operate effectively across diverse contexts, including wastewater monitoring and low-microbial-biomass clinical settings, while minimizing the impact of incomplete or low-quality genome assemblies. As a result, Seqwin provides a scalable and computationally efficient framework for microbial pathogen detection, enabling the rapid identification of genomic signatures for a wide range of downstream applications.

## Methods

Seqwin’s algorithm includes four main steps (Figure 1a). A more detailed description of each step can be found in Supplementary Note 1.

1. Generate a minimizer sketch for each input genome and build a weighted pan-genome minimizer graph (Figure 1b; a more constrained graph was described by Coombe et al. (2020) [20]).
2. Calculate a penalty score for each graph node based on the L2 norm of its absence in target genomes and presence in non-target genomes.
3. Extract connected subgraphs with average node penalty below a threshold (calculated automatically or provided by the user; Figure 1c).
4. Choose a representative sequence for each low-penalty subgraph as the signature (Figure 1d), and calculate its “conservation” (sensitivity) and “divergence” (specificity) scores by aligning the sequence to all target and non-target genomes with BLAST [10] (Supplementary Note 2).

**Figure 1.**
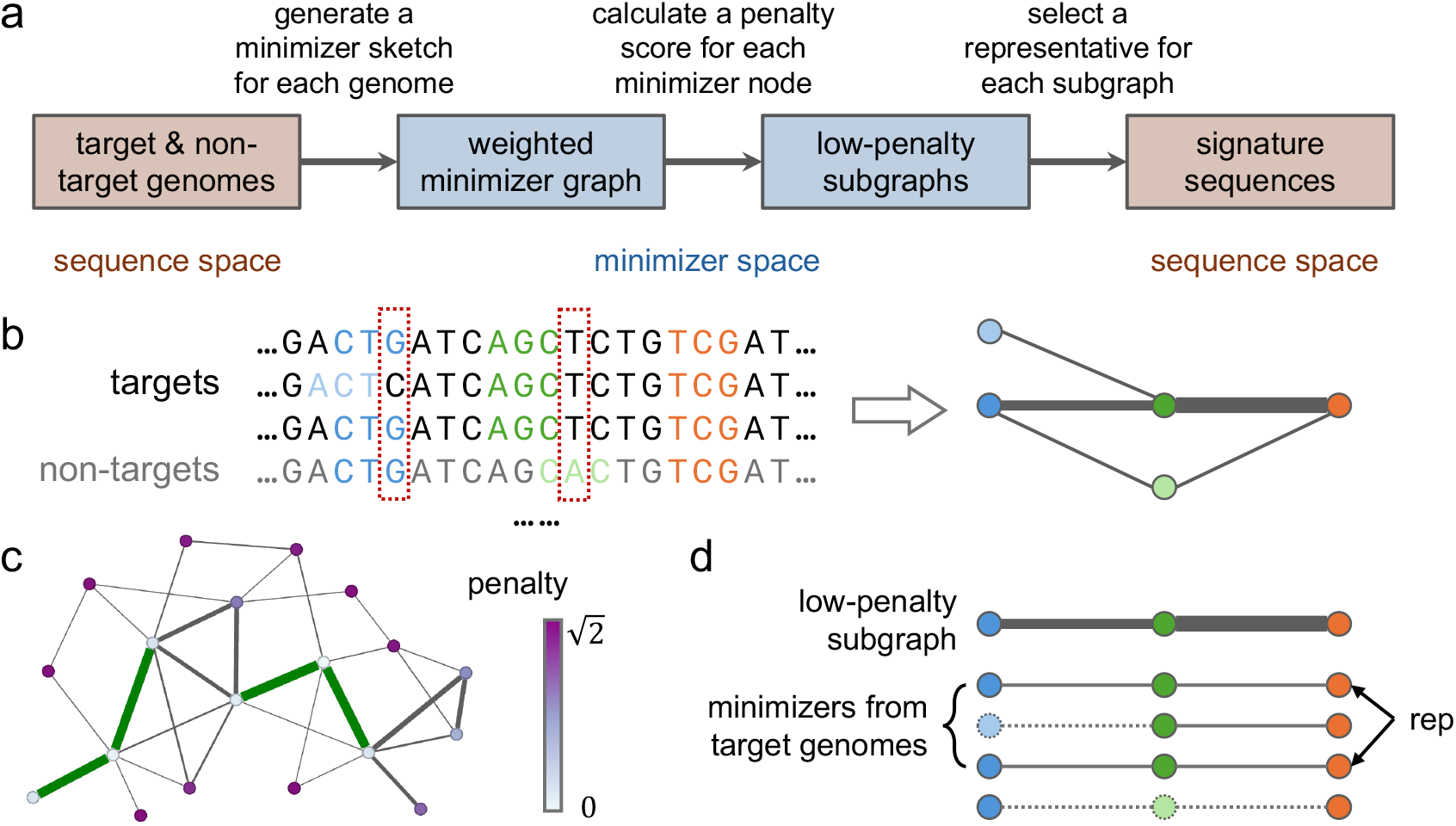
Overview of the Seqwin workflow. **a** Seqwin first compresses all input genomes into a weighted pan-genome minimizer graph. It then extracts connected low-penalty subgraphs and selects a representative for each subgraph in minimizer space. It finally converts each representative back to sequence space as the signature. **b** Graph construction (a toy example). Each unique minimizer is colored. Input genomes may yield different sets of minimizers (light blue and light orange) due to sequence variations (red dashed box). Adjacent minimizers are connected with a weighted, undirected edge in the graph. The thickness of each edge is proportional to its weight. Target genomes (black) and non-target genomes (gray) are treated equally. **c** A penalty score is calculated for each minimizer node based on the L2 norm of its absence in target genomes and presence in non-target genomes (color map). Sets of connected nodes with low average penalty and high edge weights are extracted as low-penalty subgraphs (green). **d** A representative is selected for each low-penalty subgraph. The underlying minimizers from all target genomes are listed for a low-penalty subgraph (blue, green and orange; light blue, light green and dashed lines are minimizers and edges not included in this subgraph). The most common minimizer ordering is selected as the representative (rep), and the corresponding genomic sequence becomes the signature (e.g., the first or third sequence in panel **b**).

Signatures with both high conservation and divergence are considered top candidates. Signatures with high divergence scores can still be found in non-target genomes, but only at low nucleotide identity. This reduces the chance of picking up mobile genetic elements (MGEs), which have been shown to be highly problematic as signature sequences [23] (see Supplementary Notes 1 and 2 for details).

### Generation of minimizer sketches

Seqwin first computes minimizers [24] for all input genomes with btllib [25] (version 1.7.3, with *k*-mer length *k* = 21 and window size *w* = 200 by default), including target genomes and non-target genomes. Each genome may contain one or more contigs (sequences) of unknown orientation (strand). For each sequence in a genome, a set of minimizers is sampled from its *k*-mers to form the minimizer sketch of that sequence.

We set *k* = 21 by default to lower the collision rate for reliable signature detection, which is consistent with common genome sketching defaults for assembled bacterial genomes [26]. For viral genomes, a lower *k* might be preferable. The default window size (*w* = 200) is selected to maximize the number of signatures longer than 200 bp, which is sufficient for the design of most PCR-based assays (typical amplicon length ∼100 bp). For a random minimizer sketch, the expected minimizer density is 2*/*(*w* + 1) under standard assumptions (e.g., *k* ≪ *w*) [27], implying an expected spacing of (*w* + 1)*/*2 positions between consecutive minimizers and an expected span of *w* + 1 for three consecutive minimizers (i.e., from the first to the third). Therefore, requiring at least three consecutive minimizers to represent a signature of at least 200 bp motivates *w* = 200 as a reasonable default. Throughout, we treat a run of three consecutive minimizers as the minimum evidence needed to call a signature. In practice, the choice of *w* reflects a trade-off between memory and resolution: increasing *w* reduces minimizer density and memory usage but coarsens the resolution for detecting signatures, whereas decreasing *w* increases density (and memory usage) and enables detection of shorter signatures, at the cost of representing long signatures with more consecutive minimizers and potentially reducing the number of long signatures reported.

Here, we describe the process of generating a minimizer sketch for a sequence (Algorithm S1). First, the canonical hash value of each *k*-mer in the sequence is calculated (Equation (1)) [27]. Formally,

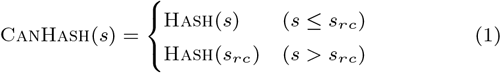

where *s* is the *k*-mer sequence, *s*_*rc*_ is the reverse complement of *s*, Hash is a hash function, and *k*-mer comparison is based on lexicographical order. Hereafter, *k*-mers and minimizers are represented by their canonical hash values unless stated otherwise. Next, for each window of *w* consecutive *k*-mers, the location of the *k*-mer with the smallest canonical hash value is selected, breaking ties by preferring the leftmost *k*-mer. The *k*-mer at this location is the minimizer of this window. The minimizer sketch of the sequence is the union of the selected *k*-mers and their locations across all windows. It is crucial that each minimizer is paired with its location in the sequence, since the same *k*-mer might be found at different locations. The minimizer sketch of a genome is the union of the sketches of each of its sequences. The minimizer sketch could also be replaced with other *k*-mer sketching methods [28, 29], as long as the “local guarantee” holds [27].

The algorithms described in this manuscript (Algorithm 1 and Algorithms S1 to S5) are not necessarily the most efficient implementations, but they have the same behavior as the Seqwin source code (version 0.2.3).

### Construction of a weighted pan-genome minimizer graph

This process is described in Algorithm 1. For each genome, we construct an undirected graph based on its minimizers, where each node corresponds to a unique minimizer (identified by its canonical hash value), and an edge is added between two nodes if the minimizers are adjacent (regardless of their ordering) in one of the genome’s sequences.

#### Algorithm 1

Construction of a weighted pan-genome minimizer graph

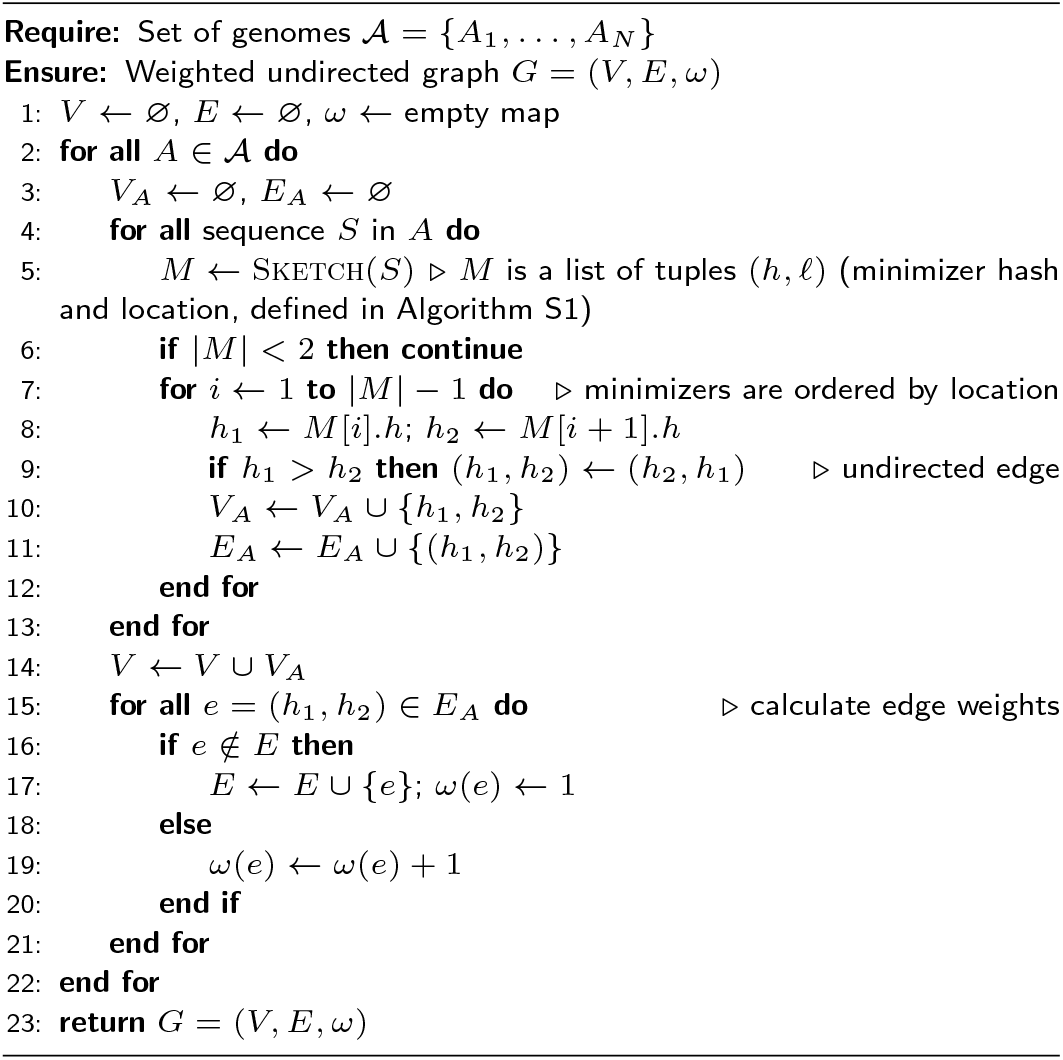

Next, the individual genome graphs are merged into a single pan-genome minimizer graph. In this merged graph, nodes represent distinct minimizers observed in any genome, and an undirected edge connects two minimizers if they are found adjacent in at least one genome. The edges are assigned a weight equal to the number of different genomes in which that minimizer adjacency occurs. In other words, if two specific minimizers appear consecutively (no matter the ordering) in the sequences of multiple genomes, the edge between their nodes is given a higher weight (reflecting the number of genomes supporting that adjacency). This results in a unified weighted graph capturing adjacency relationships of minimizers across all genomes.

### Calculation of node penalty

Since each node in the pan-genome graph represents a distinct minimizer observed in one or more genomes, suppose a minimizer node *h* is found in F_t_(*h*) target genomes and F_n_(*h*) non-target genomes. Let

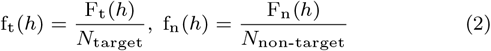

where *N*_target_ is the total number of target genomes, and *N*_non-target_ is the total number of non-target genomes. The penalty of *h* is defined as

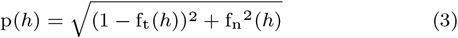

which is the L2 norm (Euclidean norm) of its absence in targets: 1 − f_t_(*h*), and presence in non-targets: f_n_(*h*). Penalty ranges from 0 (best case: the minimizer is present in all target genomes and in no non-target genomes) to 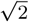 (worst case: the minimizer is absent from all targets and present in all non-targets). Thus, a lower penalty indicates that a minimizer is more sensitive and specific to the target group.

A node penalty threshold *τ*_*v*_ is used in downstream processes, and the output signatures of Seqwin are mostly derived from low-penalty nodes (p(*h*) ≤ *τ*_*v*_). *τ*_*v*_ can be determined by the user or automatically computed by Seqwin, as described next.

### Calculation of penalty threshold

Intuitively, *τ*_*v*_ should be determined by the expected *k*-mer absence and presence in target and non-target genomes, respectively. Consider a random *k*-mer *h* (not necessarily a minimizer) sampled from a random target genome (select the target genome first and then select the *k*-mer). 1 − f_t_(*h*) is the fraction of target genomes that do not include *h* (absence), and f_n_(*h*) is the fraction of non-target genomes that include *h* (presence), as defined in Equation (2). *τ*_*v*_ is then calculated with the expectations of these fractions

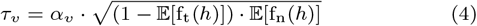

which is the geometric mean of expected *k*-mer absence and presence, multiplied by a constant *α*_*v*_. In practice, *α*_*v*_ defaults to 0.5 and can be tuned by the user to trade off the number of output signatures against their quality (e.g., increase *α*_*v*_ when more signatures are needed). We use the geometric mean so that *τ*_*v*_ will bias toward the smaller value of the two terms, resulting in a more stringent threshold.

Consider genomes as sets of *k*-mers (duplicated *k*-mers in each genome are ignored). Then these expectations can be calculated from pairwise Jaccard indices (a proof can be found in Supplementary Note 3). Since the Jaccard indices can be effectively estimated with MinHash sketches using Mash [26], *τ*_*v*_ can be calculated by running Mash on all input genomes.

Another way to estimate the expected values is to use minimizer sketches instead of MinHash sketches. This can be faster because Seqwin already generates minimizer sketches for all input genomes. As previously reported, minimizer sketches on average underestimate the fraction of shared *k*-mers between two sequences [30], resulting in biased estimates of the expectations. However, since this bias leads to overestimation of *k*-mer absence in targets (1 − E[f_t_(*h*)]) and underestimation of *k*-mer presence in non-targets (E[f_n_(*h*)]), the bias of the resulting geometric mean is smaller (Equation (4)). Consequently, this provides a practical solution for estimating *τ*_*v*_ defined in Equation (4), particularly when processing large datasets where calculating pairwise Jaccard indices is computationally prohibitive. Seqwin implements this method as a faster alternative for calculating *τ*_*v*_, along with the unbiased Mash implementation.

### Extraction of low-penalty subgraphs

The minimizer graph is filtered to remove edges with negligible weights and isolated nodes (Supplementary Note 4). Disjoint low-penalty subgraphs are then extracted from the filtered graph (Algorithm S2). Each subgraph is a set of connected minimizer nodes whose average node penalty (Equation (3)) does not exceed the threshold *τ*_*v*_. The procedure first identifies all candidate seed nodes with penalty ≤ *τ*_*v*_. Starting from each seed, a greedy breadth-first search (BFS) expansion is performed by iteratively adding the adjacent node with the lowest penalty, as long as including that node keeps the subgraph’s average penalty below *τ*_*v*_. This expansion continues until no more neighboring nodes can be added or the subgraph reaches a specified maximum size (100 by default). Subgraphs meeting the minimum size requirement (3 by default) are retained, and their nodes are marked used so subgraphs remain disjoint (no node is part of more than one subgraph). Seeds are processed in random order, and the final list of subgraphs is shuffled to balance subgraph sizes in downstream processes (subgraphs created first tend to be larger due to the greedy optimization).

### Choosing a representative sequence for each low-penalty subgraph

For each low-penalty subgraph (a set of connected minimizer nodes from the graph), a representative minimizer ordering is determined and the corresponding genomic sequence is output as a candidate signature. The procedure involves 1) finding the maximal consecutive occurrence of the subgraph’s minimizers in each target genome, and 2) choosing the most common one across all target genomes, weighted by length (Algorithm S3 and Figure 1d). A more detailed illustration of this process can be found in Figure S1.

In each target genome that contains one or more minimizers from a given subgraph, we identify the longest segment of those minimizers that appear consecutively in the genome’s sequence (allowing for at most one intervening minimizer not in the subgraph). Note that there could be repetitive segments in a single genome. Two minimizers are defined as consecutive if 1) they appear in the same sequence of that genome, 2) the difference between their indices in the minimizer sketch is less than or equal to 2. This yields, for each genome, an ordered tuple of minimizer hashes representing that subgraph’s segment in the genome.

Among all target genomes, the most common minimizer ordering (treating forward and reverse ordering as equivalent) is then selected as the subgraph’s representative minimizer ordering. Prevalence is measured by the number of target genomes in which a given ordering occurs, weighted by the length (number of minimizers) of the ordering to favor longer sequences. The orientation of the representative ordering is chosen to match the strand orientation that is more commonly observed for that sequence in the target genomes. One of the genomes supporting this representative minimizer ordering is used to determine its actual genomic sequence (the representative sequence), based on the minimizer coordinates in that genome. Representative sequences that satisfy user-defined length thresholds (≥ 200 bp by default) are output as candidate signatures. Candidates are evaluated for their sensitivity and specificity, measured as conservation and divergence, respectively (Supplementary Note 2).

## Results

### Seqwin outperforms existing tools in signature quality, running time and memory usage

To test Seqwin’s ability to identify sensitive and specific signatures, we compared it with other tools. We selected Fur as a fast, memory-efficient method and Unikseq and Neptune as representative *k*-mer assembly methods (Table 1). Note, NAUniSeq was not included in the benchmark given we were unable to successfully run it on our evaluation dataset due to missing files that caused installation errors. We first used the dataset published with Fur [11] to benchmark the four tools (Seqwin, Fur, Unikseq and Neptune). The Fur dataset consists of 33 *Escherichia coli* genomes from 6 different strains, labeled A (6 genomes), B1 (14 genomes), B2 (5 genomes), D (2 genomes), E (4 genomes) and F (2 genomes). For each run of each tool, genomes under one of the strains were used as targets (e.g., strain A), and genomes under all other strains (e.g., B1, B2, D, E and F) were used as non-targets. Thus, each tool generated 6 different sets of signatures, summarized in Table S1. Each signature was evaluated for its sensitivity (conservation) and specificity (divergence), with results shown in Figure S2. As a supplement to the divergence score, for each signature we also counted the fraction of non-target genomes with a BLAST hit, shown in Figure S3.

Next, we benchmarked all four tools against microbial genomes retrieved from NCBI Taxonomy. We downloaded all available genomes (August 2025) under three pathogenic taxa and their neighboring taxa: *Clostridioides difficile* (ID 1496, 3,995 genomes), *Mycobacterium tuberculosis* (ID 1773, 8,296 genomes) and *Salmonella enterica* subspecies *enterica* (ID 59201, 14,822 genomes). Here, “neighboring taxa” are those that diverge from a common ancestral node into separate sister lineages. For example, as the only species in the genus *Clostridioides, C. difficile*’s neighbors encompass all other genera under the shared family, *Peptostreptococcaceae*. Genomes from all assembly quality levels were included, while those labeled as “Atypical genomes” and “Genomes from large multi-isolate projects” were excluded. Genomes under each pathogenic taxon were used as targets and genomes under the neighboring taxa were used as non-targets. For each dataset, we sampled 100 or 1,000 genomes as input (targets and non-targets combined), with results shown in Table 2 and Figures 2 and S4. To retain genome diversity, we included at least 20 non-target genomes for each of the 100-genome datasets, since target genomes far outnumbered non-target genomes in the full datasets (Table 2). Metadata of all NCBI genomes can be found in Supplementary Data 2-4.

**Table 2.** Benchmarking of Seqwin, Fur, Unikseq, and Neptune on genomes from NCBI Taxonomy.

| Target taxon | # genomes<sup>1</sup> | Tool | # signatures | Median length (bp) | Median conservation | Median divergence | Wall-clock time (s)<sup>2</sup> | Peak memory usage (GB)<sup>3</sup> |
| --- | --- | --- | --- | --- | --- | --- | --- | --- |
| *C. difficile*<br>(ID 1496) | 100 (20) | Seqwin | 33 | 271 | 0.994 | 0.12 | 12.2 | 3.42 |
|  |  | Fur | 0 | N/A | N/A | N/A | 10.8 | 3.54 |
|  |  | Unikseq | 9,074 | 237 | 0.975 | 0.067 | 1,090 | 78.8 |
|  |  | Neptune | 3,679 | 327 | 0.962 | 0.025 | 2,450 | 13.5 |
|  | 1,000 (167) | Seqwin | 227 | 313 | 0.992 | 0.13 | 34.5 | 5.14 |
|  |  | Fur | 0 | N/A | N/A | N/A | 68.1 | 4.21 |
|  |  | Unikseq | 10,989 | 186 | 0.982 | 0.032 | 10,600 | 565 |
|  |  | Neptune | 4,596 | 212 | 0.977 | 0.014 | 18,700 | 13.9 |
|  | 3,995 (667) | Seqwin | 156 | 312 | 0.993 | 0.13 | 70.0<sup>4</sup> | 7.60 |
| *M. tuberculosis*<br>(ID 1773) | 100 (20) | Seqwin | 1,674 | 1184 | 0.994 | 0.14 | 18.3 | 4.29 |
|  |  | Fur | 655 | 372 | 0.995 | 0.0087 | 12.4 | 6.14 |
|  |  | Unikseq | 10,520 | 199 | 1.00 | 0.16 | 1,130 | 71.8 |
|  |  | Neptune | 1,843 | 350 | 0.992 | 0.039 | 1,360 | 13.5 |
|  | 1,000 (77) | Seqwin | 150 | 312 | 0.997 | 0.090 | 40.0 | 5.32 |
|  |  | Fur | 0 | N/A | N/A | N/A | 63.2 | 7.46 |
|  |  | Unikseq | 103 | 172 | 0.993 | 0.020 | 11,700 | 576 |
|  |  | Neptune | 30 | 301 | 0.993 | 0.0086 | 2,760 | 14.8 |
|  | 8,296 (637) | Seqwin | 208 | 350 | 0.998 | 0.091 | 194<sup>4</sup> | 11.7 |
| *S. enterica*<br>subsp. *enterica*<br>(ID 59201) | 100 (20) | Seqwin | 382 | 333 | 0.990 | 0.048 | 13.7 | 3.74 |
|  |  | Fur | 0 | N/A | N/A | N/A | 11.4 | 6.22 |
|  |  | Unikseq | 593 | 249 | 0.339 | 0.017 | 1,260 | 77.2 |
|  |  | Neptune | 24 | 800 | 0.685 | 0.0036 | 206 | 14.7 |
|  | 1,000 (21) | Seqwin | 319 | 317 | 0.990 | 0.053 | 40.0 | 5.39 |
|  |  | Fur | 0 | N/A | N/A | N/A | 52.9 | 4.91 |
|  |  | Unikseq | 542 | 233 | 0.337 | 0.021 | 13,200 | 624 |
|  |  | Neptune | 23 | 382 | 0.797 | 0.0029 | 2,770 | 15.6 |
|  | 14,822 (304) | Seqwin | 275 | 321 | 0.990 | 0.049 | 313<sup>4</sup> | 22.3 |

<sup>1</sup> Including neighboring non-target genomes. The number of non-target genomes is shown in parentheses. Percentage of complete genomes (assembly level at or above chromosome) in each full dataset: 7.43% (*C. difficile*), 12.03% (*M. tuberculosis*), 13.73% (*S. enterica*). Metadata of all genomes can be found in Supplementary Data 2-4.
<sup>2</sup> 20 CPU threads/processes were used for Seqwin, Fur and Neptune. Unikseq only supported a single CPU thread.
<sup>3</sup> Neptune stores *k*-mers of input genomes as temporary files on disk.
<sup>4</sup> When Seqwin was run on the full datasets, minimizer sketches were used to calculate the penalty thresholds to reduce running time (see Methods and Supplementary Note 5).

**Figure 2.**
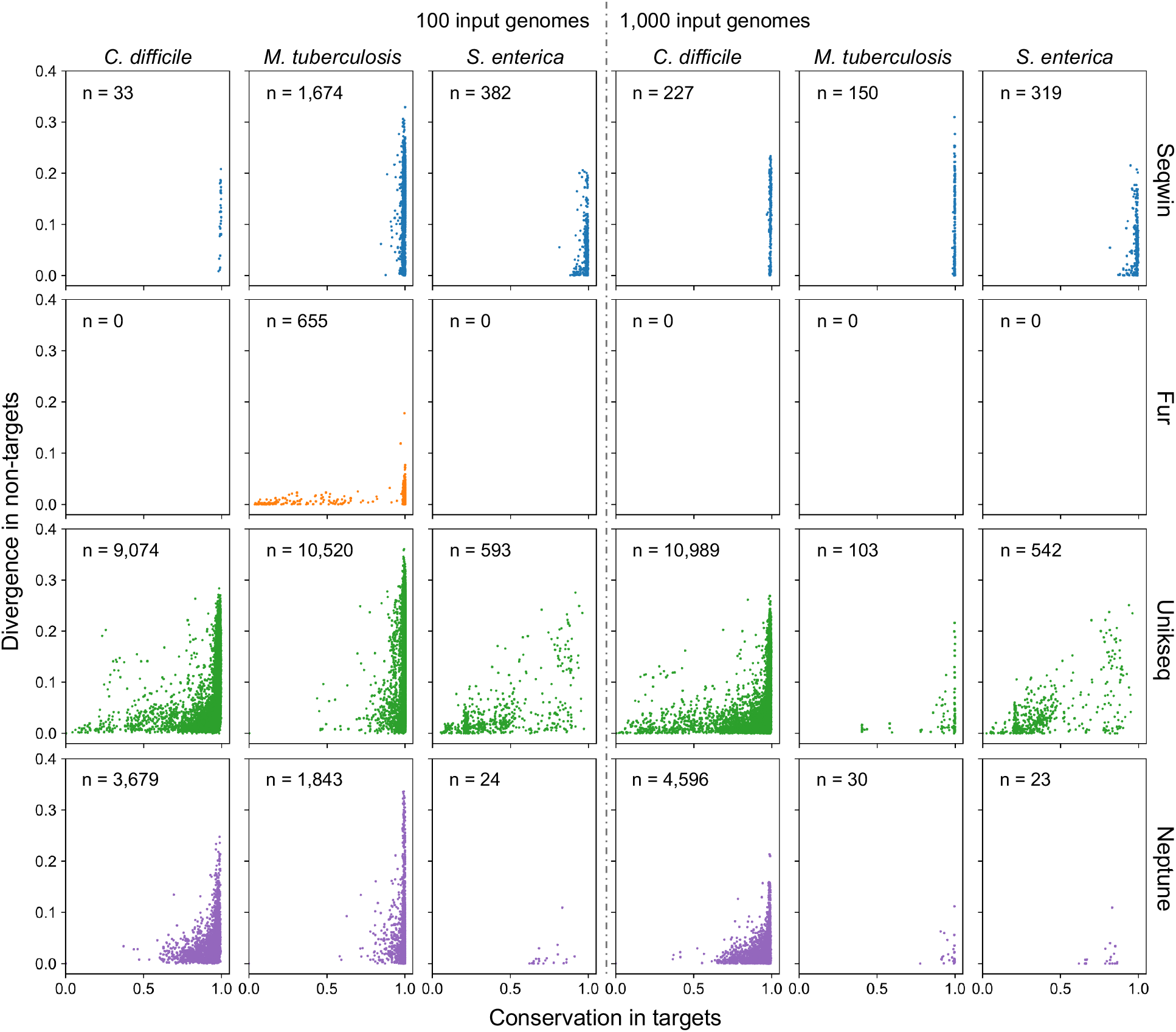
Benchmarking of Seqwin, Fur, Unikseq and Neptune using genomes downloaded from NCBI Taxonomy [31]. Each data point represents a signature sequence, generated by one of the tools using genomes under a pathogenic taxon as targets (e.g., *C. difficile*), and genomes under its neighboring taxa as non-targets. Blue, orange, green and purple represent Seqwin, Fur, Unikseq and Neptune, respectively. The number of output signatures (data points) in each setting is shown in each scatter plot. 100 or 1,000 genomes are sampled from each NCBI full dataset (Table 2), and used as inputs to the tools. See Supplementary Note 2 for the calculation of conservation and divergence.

Compared to Fur, Seqwin identified more signatures with comparable running time and peak memory (Tables S1 and 2). Fur produced zero signatures in several experiments, most likely due to its stringent search strategy. Compared to Unikseq, Seqwin’s output signatures had similar median lengths (Tables S1 and 2) in most settings. Although Unikseq produced more signatures, many of them had lower conservation scores (Figures S2 and 2). This was especially true for *S. enterica* (Table 2 and Figure 2), where Unikseq identified only a handful of high-conservation signature sequences. Seqwin was also more efficient with respect to wall-clock time and peak memory usage. Neptune’s output signatures resembled those of Unikseq, with higher conservation but still less than Seqwin (Table 2 and Figure 2). It also required more running time and peak memory than Seqwin (Tables S1 and 2). Note that Neptune stores *k*-mers of input genomes as temporary files on disk, which reduces peak memory usage but can increase runtime.

Lists of genomes used in all experiments can be found in Supplementary Data 1-4. Benchmark details can be found in Supplementary Note 5.

### Seqwin efficiently scales up to thousands of bacterial genomes

Next, we evaluated only Seqwin on all genomes downloaded from NCBI Taxonomy, as Fur did not generate any signatures for the benchmark settings with 1,000 genomes, and Unikseq was estimated to require terabytes of memory (Table 2). For nearly 15,000 *S. enterica* genomes, Seqwin finished in only 5 minutes using 20 CPU cores and 22 GB peak memory (Table 2). Seqwin also maintained similar signature quantity and quality to those obtained with 1,000 genomes, as shown in Figure 3. Note that the penalty thresholds in these experiments were calculated with minimizer sketches to reduce running time (see Methods for more details).

**Figure 3.**
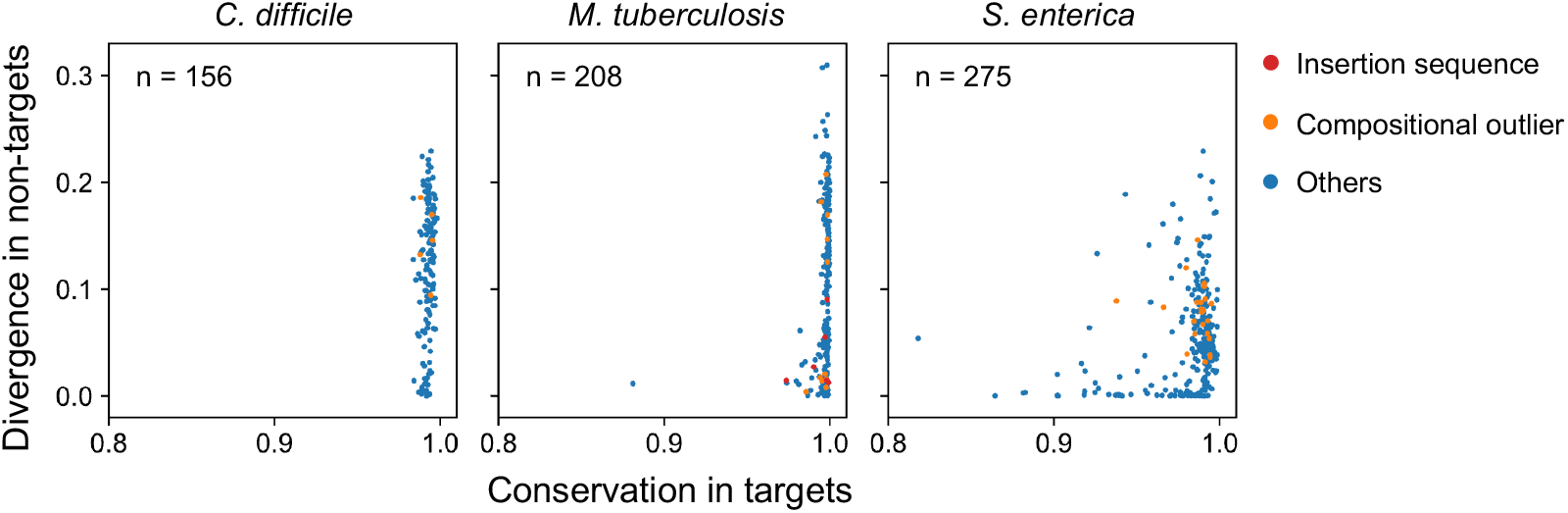
Output signatures of Seqwin using all genomes downloaded from NCBI Taxonomy. Each data point represents a signature sequence, generated by Seqwin using genomes under a pathogenic taxon as targets (e.g., *C. difficile*), and genomes under its neighboring taxa as non-targets. The number of output signatures (data points) in each setting is shown in each scatter plot. Signatures overlapping with potential MGEs are labeled in red and orange, all others are blue. Number of orange data points (left to right): 5, 11 and 23. Number of red data points (left to right): 0, 6 and 0.

In addition, we annotated the reference genomes of the target pathogens and identified genes and mobile genetic elements (MGEs), using eggNOG-mapper [32] and the Mobilome Annotation Pipeline (MAP) under MGnify [33], respectively. “Compositional outliers” were potential MGEs identified by MAP (Figure 3), indicating genomic regions with abnormal composition (e.g., GC content) compared to their contexts. Insertion sequences were identified by MAP and were also confirmed by gene annotations, with gene products annotated as transposase, integrase or resolvase. In all three experiment settings, less than 10% of the signatures overlapped with predicted MGEs (compositional outliers and insertion sequences). Moreover, only *M. tuberculosis* had signatures overlapping with insertion sequences (Figure 3), and their median divergence was 0.021, which is lower than the median divergence of all *M. tuberculosis* signatures (Table 2). Annotation results of all signatures shown in Figure 3 can be found in Supplementary Data 5-7. Details of the annotation process can be found in Supplementary Note 6.

## Discussion

We demonstrate that Seqwin represents a significant advance in scalable microbial genome signature discovery through benchmarking experiments. Seqwin identified genomic signatures in tens of thousands of genomes, using target and non-target genome sets as input. By coupling a minimizer-graph strategy with tolerance for sequence variation and genomic diversity, Seqwin bypasses strict search criteria and high memory usage, critical limitations of previous methods. Through experimental evaluation, we show that Seqwin consistently achieves higher signature sensitivity and specificity across both small and large-scale datasets while using fewer computational resources than other approaches. Although wet-lab validation was not performed in this study (planned as future work), we envision Seqwin being coupled with established primer and probe design software, such as varVAMP [34], Olivar [35] and PrimalScheme [36], supporting sensitive, specific PCR assay design for clinical and public health surveillance.

However, there are two major open problems we leave for future work. First, Seqwin estimates specificity from the provided non-target genomes and considers all non-targets equally. Consequently, specificity is best estimated when the non-target set is restricted to close phylogenetic neighbors, ideally using well-curated and high-quality genomes. Although Seqwin is flexible and tolerates some noise in the non-target set, including a large number of very distantly related non-target genomes can bias specificity estimation. In practice, it is suggested to perform an additional screening of candidate signatures against a comprehensive background and examine hits outside the intended search space. Taxonomic mislabelling in public databases is another challenge that Seqwin itself cannot resolve. Using a genome-based taxonomy database such as GTDB [37] may alleviate this issue. A potential solution is to weight non-target genomes by sequence similarity, for example using Jaccard indices, so that specificity estimation is driven more strongly by the most relevant near-neighbor genomes. Second, Seqwin is designed to identify single genomic regions that distinguish the target group from the non-target group, rather than combinations of multiple regions that are informative only when considered together. As a result, Seqwin may miss combinatorial signatures in which no individual region is sufficiently conserved across the full target set, even though the union of those regions could still distinguish targets from non-targets. This limitation is relevant when designing assays for antimicrobial-resistant (AMR) strains of a pathogen, as AMR can depend on combinations of mutations or genes across multiple loci rather than on a single diagnostic region [38]. Support for identifying combinatorial signatures is under development and is planned for a future Seqwin update.

In summary, inspired by approaches pioneered by Insignia two decades ago, Seqwin represents a highly sensitive and discriminative computational approach for microbial genome signature discovery. We anticipate Seqwin will enable signature discovery at unprecedented scale and support automated, sensitive, specific PCR assay design for clinical and environmental applications.

## Data availability

NCBI accessions and metadata of all genomes used in this study can be found in Supplementary Data. Benchmarking datasets, outputs, and scripts are available on Zenodo https://doi.org/10.5281/zenodo.19176444.

## Author contributions statement

M.X.W., L.B.S., and T.J.T. conceived the project; M.X.W. and B.K. conceived the algorithm; M.X.W. developed the code; M.X.W., M.G.N., S.Z., and T.J.T. conceived the experiments; M.X.W. conducted the experiments; M.X.W., M.G.N., and T.J.T. drafted the original manuscript; all authors reviewed the manuscript.

## Acknowledgments

The authors thank Dr. Adam Phillippy and Dr. Jingjing Wu for valuable feedback and suggestions.

