## Supplementary Information for "Seqwin: Ultrafast identification of signature sequences in microbial genomes"

### (Supplementary Information) Seqwin: Ultrafast identification of signature sequences in microbial genomes

Michael X. Wang et al.

Table S1: Benchmarking of Seqwin, Fur, Unikseq, and Neptune on the Fur dataset (33 *E. coli* genomes)

| Target strain | # genomes[1] | Tool | # signatures | Median length (bp) | Wall-clock time (s)[2] | Peak memory usage (GB)[3] |
|---|---|---|---|---|---|---|
| A | 6 | Seqwin | 98 | 297 | 25.6 | 0.770 |
| | | Fur | 0[4] | 0 | 48.9 | 0.729 |
| | | Unikseq | 303 | 231 | 456 | 32.6 |
| | | Neptune | 25 | 397 | 957 | 0.150 |
| B1 | 14 | Seqwin | 98 | 323 | 23.4 | 0.585 |
| | | Fur | 0[4] | 0 | 35.6 | 0.570 |
| | | Unikseq | 391 | 192 | 463 | 32.6 |
| | | Neptune | 17 | 433 | 954 | 0.151 |
| B2 | 5 | Seqwin | 633 | 324 | 30.4 | 0.776 |
| | | Fur | 25 | 499 | 59.5 | 0.738 |
| | | Unikseq | 442 | 179 | 457 | 32.1 |
| | | Neptune | 109 | 559 | 998 | 0.147 |
| D | 2 | Seqwin | 583 | 332 | 33.0 | 0.967 |
| | | Fur | 31 | 884 | 62.9 | 0.773 |
| | | Unikseq | 236 | 213 | 463 | 32.4 |
| | | Neptune | 93 | 953 | 1010 | 0.151 |
| E | 4 | Seqwin | 201 | 324 | 26.1 | 0.771 |
| | | Fur | 18 | 466 | 53.0 | 0.747 |
| | | Unikseq | 198 | 145 | 455 | 32.0 |
| | | Neptune | 58 | 282 | 978 | 0.152 |
| F | 2 | Seqwin | 485 | 344 | 31.8 | 0.964 |
| | | Fur | 17 | 692 | 62.1 | 0.770 |
| | | Unikseq | 267 | 223 | 457 | 32.3 |
| | | Neptune | 85 | 650 | 995 | 0.154 |

[1] Genomes from all other strains were used as non-targets. Genome accessions can be found in Supplementary Data 1.
[2] A single CPU thread was used for all tools.
[3] Neptune stores $k$-mers of input genomes as temporary files on disk.
[4] In the original Fur paper, Fur output 0 and 0.1 kb of signatures for strain A and B1, respectively.

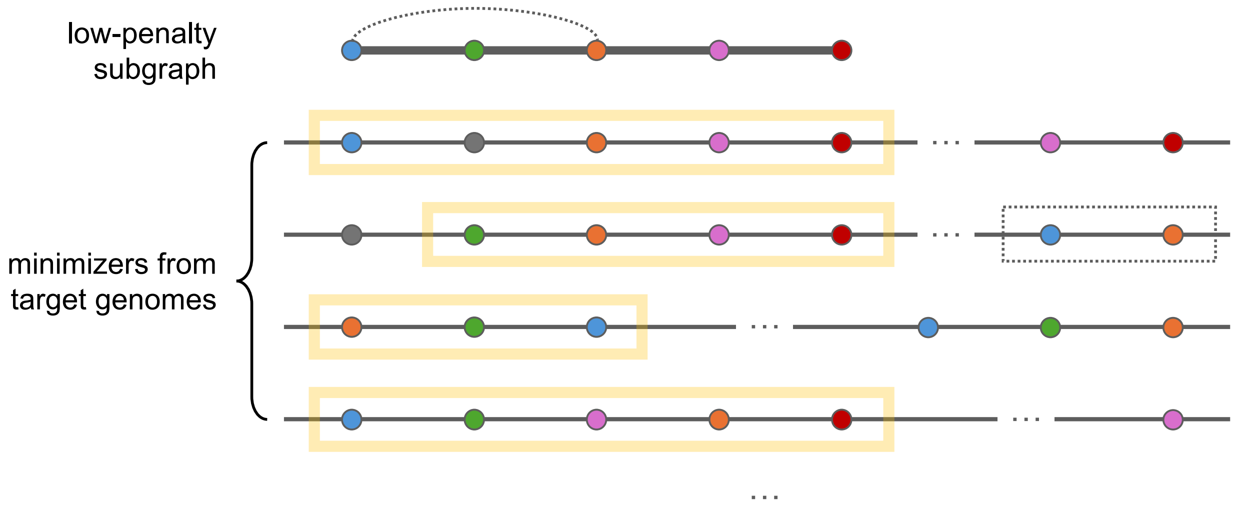


Figure S1: This figure supplements Figure 1d by illustrating how the representative minimizer ordering is chosen for a low-penalty subgraph. Each unique minimizer is shown as a colored dot, and minimizers not included in the subgraph are shown as gray dots.
**(1)** In the first target genome, since the blue and orange minimizers have only one intervening minimizer (gray), they are still considered consecutive. Thus, the five minimizers on the left constitute the maximal consecutive occurrence of the subgraph's minimizers, or simply the **maximal occurrence** (light yellow box).
**(2)** In the second genome, the blue minimizer is missing, yet the other four minimizers still appear as consecutive. Note that the blue and orange minimizers also appear to be consecutive in another part of the genome, resulting in a low-weight edge in the graph that is later pruned (dashed box and line).
**(3)** In the third genome, the three colored minimizers appear consecutively twice, and the minimizers in the first occurrence are in reverse order.
**(4)** In the fourth genome, all five colored minimizers appear consecutively, but the order of the orange and pink minimizers is reversed.
The most common **maximal occurrence** in all target genomes (regardless of orientation, weighted by length) is the representative minimizer ordering.

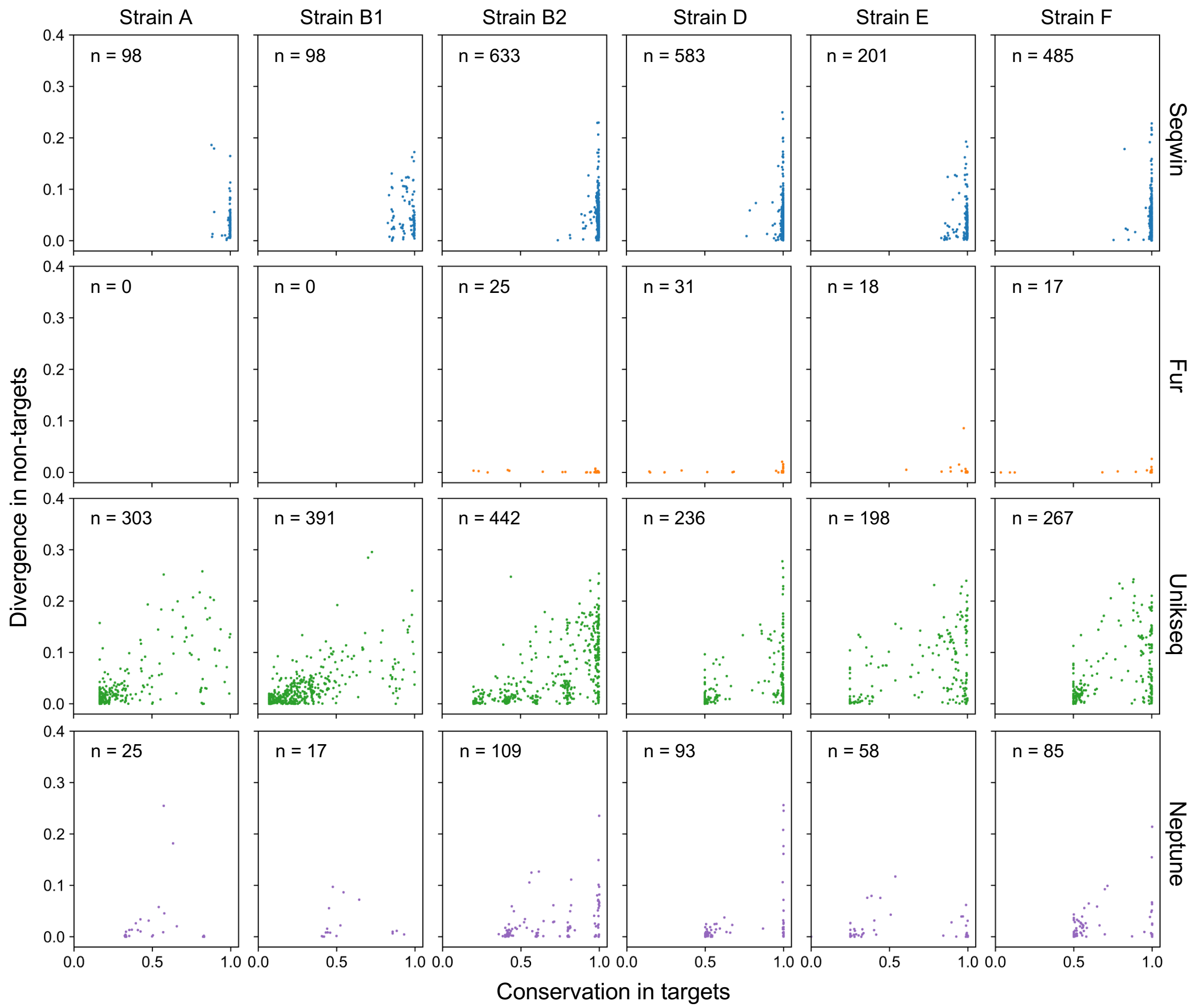


Figure S2: Benchmarking of Seqwin, Fur, Unikseq, and Neptune using the Fur dataset (33 *E. coli* genomes). Each data point represents a signature sequence, generated by one of the tools using genomes under one strain as targets, and genomes under the other five strains as non-targets (e.g., plots in column "strain A" are generated using genomes under strain A as targets). Blue, orange, green and purple represent Seqwin, Fur, Unikseq, and Neptune, respectively. The number of output signatures (data points) in each setting is shown in each scatter plot. See Supplementary Note 2 for the calculation of conservation and divergence.

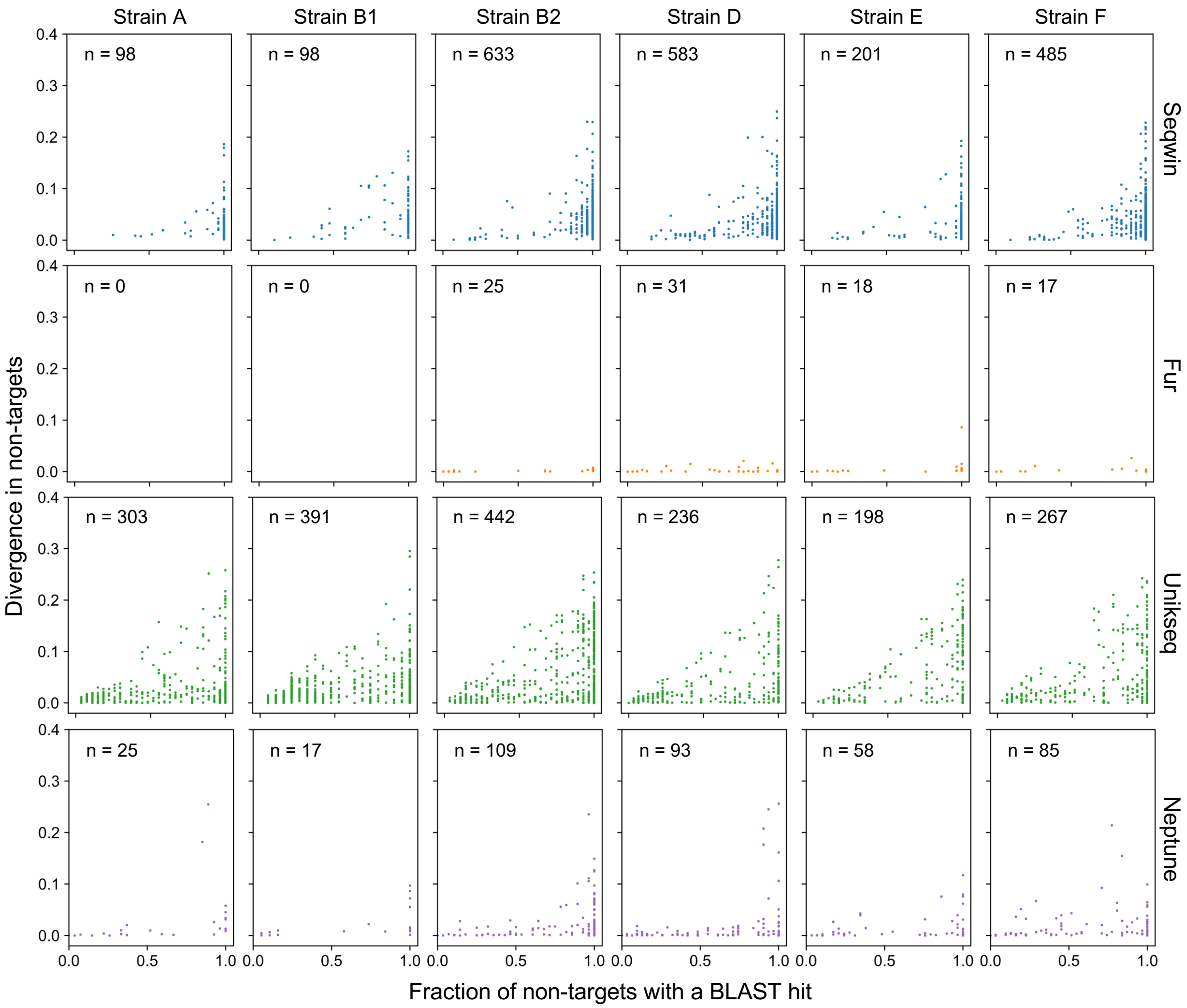


Figure S3: This figure was produced with the same sets of signatures as Figure S2, using the same layout. For each signature, the fraction of non-target genomes with at least one BLAST hit is shown (x-axis). For example, if a signature is found in half of the non-targets, the fraction would be 0.5. If a signature is found in none of the non-targets (fraction is 0), its divergence would also be 0. For a signature found in all non-target genomes and a divergence of 0.2, it means that on average 20% of the nucleotides are different in the corresponding non-target regions. See Supplementary Note 2 for more details.

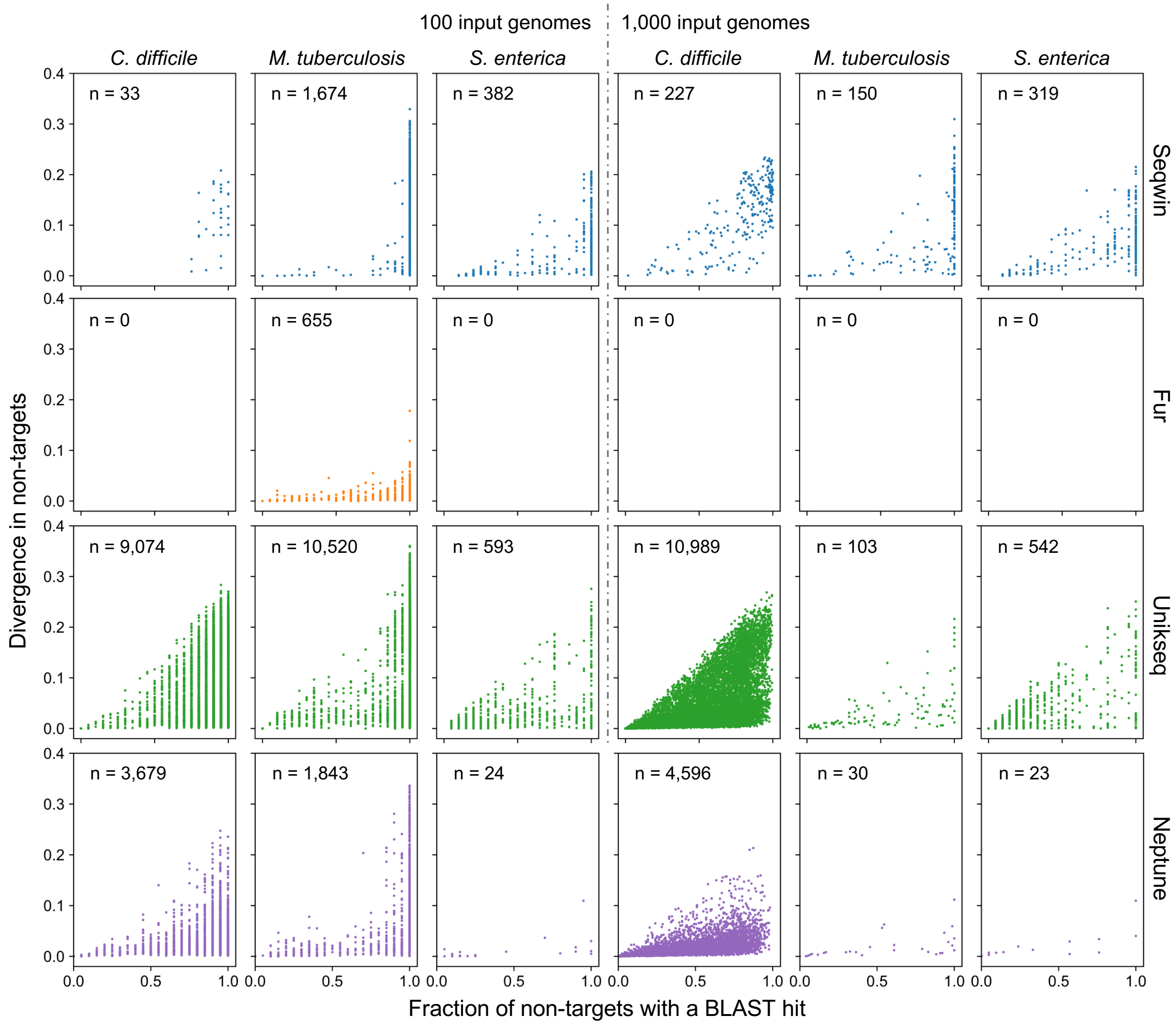


Figure S4: This figure was produced with the same sets of signatures as Figure 2, using the same layout. For each signature, the fraction of non-target genomes with at least one BLAST hit is shown (x-axis). For example, if a signature is found in half of the non-targets, the fraction would be 0.5. If a signature is found in none of the non-targets (fraction is 0), its divergence would also be 0. For a signature found in all non-target genomes and a divergence of 0.2, it means that on average 20% of the nucleotides are different in the corresponding non-target regions. See Supplementary Note 2 for more details.

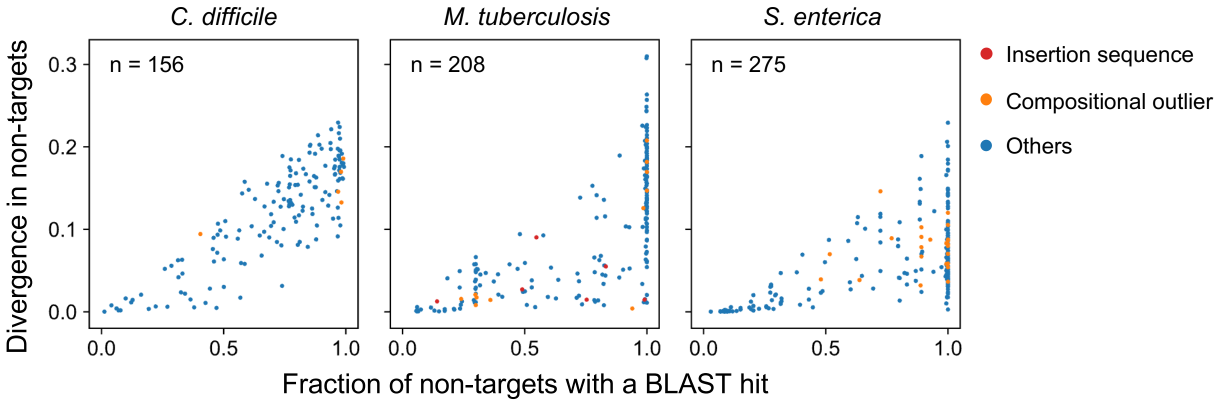


Figure S5: This figure was produced with the same sets of signatures as Figure 3, using the same layout. For each signature, the fraction of non-target genomes with at least one BLAST hit is shown (x-axis). For example, if a signature is found in half of the non-targets, the fraction would be 0.5. If a signature is found in none of the non-targets (fraction is 0), its divergence would also be 0. For a signature found in all non-target genomes and a divergence of 0.2, it means that on average 20% of the nucleotides are different in the corresponding non-target regions. See Supplementary Note 2 for more details.

**Algorithm S1** Generation of the minimizer sketch for a sequence

**Require:** Sequence $S$, integer $k$, integer $w$
**Ensure:** Minimizer sketch $M$ as a list of tuples $(h, \ell)$ ▷ canonical hash value and location

**function** SKETCH($S$)
  **if** $|S| < w + k - 1$ **then return** [ ] ▷ too short to generate a minimizer, return an empty list
  $L \leftarrow [\,]$
  **for** $\ell \leftarrow 1$ **to** $|S| - k + 1$ **do**
    $h \leftarrow$ CANHASH( $S[\,\ell \ldots \ell + k - 1\,]$ ) ▷ canonical hash value of each $k$-mer (Equation (1))
    APPEND($L$, $(h, \ell)$) ▷ add to the end of the list
  **end for**
  $M \leftarrow [\,]$
  **for** $i \leftarrow 1$ **to** $|L| - w + 1$ **do** ▷ find the minimizer for each window
    $(h, \ell) \leftarrow \min(\,L[\,i \ldots i + w - 1\,]\,)$ ▷ lexicographical minimum (sort by hashes first, then locations)
    **if** $(h, \ell) \notin M$ **then** APPEND($M$, $(h, \ell)$) ▷ note $(h, \ell_1) \neq (h, \ell_2)$ if $\ell_1 \neq \ell_2$
  **end for**
  **return** $M$
**end function**

**Algorithm S2** Extraction of disjoint low-penalty subgraphs via greedy BFS expansion

**Require:** Filtered graph $G = (V, E, p)$ with node penalties $\mathrm{p}(h)$ for each $h \in V$
**Require:** Node penalty threshold $\tau_v$; subgraph size bounds $(n_{\min}, n_{\max})$
**Ensure:** Set $\mathcal{V}$ as disjoint connected subsets of $V$

seeds $\leftarrow \{\, h \in V : \mathrm{p}(h) \leq \tau_v \,\}$ ▷ seed nodes with low penalty
SHUFFLE(seeds) ▷ process seeds in random order to avoid bias
$\mathcal{V} \leftarrow \varnothing$; used $\leftarrow \varnothing$ ▷ initialize set of subgraphs and set of used nodes
**for all** $s \in$ seeds **do**
  **if** $s \in$ used **then continue** ▷ skip seed if already used in another subgraph
  $V_s \leftarrow \{s\}$; sum $\leftarrow \mathrm{p}(s)$ ▷ initialize subgraph and sum of node penalties
  $F \leftarrow \{\, h \in \text{NEIGHBORS}(s) : h \notin \text{used} \wedge h \notin V_s \,\}$ ▷ add neighboring nodes of $s$ in $G$ to frontier
  **while** $F \neq \varnothing$ **and** $|V_s| < n_{\max}$ **do**
    $x \leftarrow \arg\min_{h \in F}\, \mathrm{p}(h)$ ▷ neighbor with lowest penalty
    $F \leftarrow F \setminus \{x\}$ ▷ whether $x$ is accepted or not, remove it from frontier
    $\text{avg} \leftarrow \dfrac{\text{sum} + \mathrm{p}(x)}{|V_s| + 1}$ ▷ average penalty if $x$ is added
    **if** avg $\leq \tau_v$ **then**
      $V_s \leftarrow V_s \cup \{x\}$; sum $\leftarrow$ sum $+ \mathrm{p}(x)$
      $F \leftarrow F \cup \{\, h \in \text{NEIGHBORS}(x) : h \notin \text{used} \wedge h \notin V_s \,\}$ ▷ add new neighbors of $x$
    **end if**
  **end while**
  **if** $|V_s| \geq n_{\min}$ **then**
    $\mathcal{V} \leftarrow \mathcal{V} \cup \{V_s\}$; used $\leftarrow$ used $\cup V_s$
  **end if**
**end for**
SHUFFLE($\mathcal{V}$) ▷ shuffle subgraphs for balanced downstream processing
**return** $\mathcal{V}$

**Algorithm S3** Choosing a representative minimizer ordering for a low-penalty subgraph

**Require:** Low-penalty subgraph $V_s$ as a set of nodes (minimizers)
**Require:** Set of target genomes $\mathcal{A}_{\text{target}} = \{A_1, \ldots, A_{N_{\text{target}}}\}$
**Ensure:** $H_{\text{rep}}$ as an ordered tuple of minimizers
$\mathcal{H} \leftarrow$ empty multiset ▷ longest consecutive segment of minimizers in each genome
**for all** $A \in \mathcal{A}_{\text{target}}$ **do**
  $\mathcal{H}_A \leftarrow \varnothing$ ▷ consecutive segments in the current genome
  **for all** sequence $S$ in $A$ **do**
    $M \leftarrow \text{SKETCH}(S)$ ▷ $M$ is a list of tuples $(h, \ell)$ (defined in Algorithm S1)
    **if** $|M| = 0$ **then continue**
    $M_s \leftarrow [\,]$
    **for** $i \leftarrow 1$ **to** $|M|$ **do** ▷ get minimizers in $M$ that are also present in $V_s$
      $h \leftarrow M[i].h$ ▷ get the minimizer hash value and ignore its location
      **if** $h \in V_s$ **then** $\text{APPEND}(M_s, (i, h))$ ▷ store the index of this minimizer in $M$
    **end for**
    $\mathcal{H}_A \leftarrow \mathcal{H}_A \cup \text{CONSECUTIVESEGMENTS}(M_s)$ ▷ see Algorithm S4
  **end for**
  **if** $\mathcal{H}_A \neq \varnothing$ **then**
    $H_{\max} \leftarrow \arg\max_{H \in \mathcal{H}_A} |H|$ ▷ get the longest consecutive segment
    $\text{ADD}(\mathcal{H}, H_{\max})$ ▷ add $H_{\max}$ to the multiset (duplicates are allowed)
  **end if**
**end for**
$H_{\text{rep}} \leftarrow \text{MOSTCOMMON}(\mathcal{H})$ ▷ see Algorithm S5
**return** $H_{\text{rep}}$

**Algorithm S4** Find segments of consecutive minimizers

**Require:** $M_s$ as a list of tuples $(i, h)$, where $h$ is the hash value of a minimizer and $i$ is its index
**Ensure:** $\mathcal{H}_s$ as a set of tuples; each tuple consists of hashes with consecutive indices
**function** $\text{CONSECUTIVESEGMENTS}(M_s)$
  $\mathcal{H}_s \leftarrow \varnothing$
  **if** $|M_s| = 0$ **then return** $\mathcal{H}_s$
  $(i, h) \leftarrow M_s[1]$
  $H \leftarrow [\,h\,]$; prev $\leftarrow i$ ▷ initialize the first consecutive segment
  **if** $|M_s| = 1$ **then return** $\{\text{TUPLE}(H)\}$ ▷ convert list to tuple
  **for** $j \leftarrow 2$ **to** $|M_s|$ **do**
    $(i, h) \leftarrow M_s[j]$
    **if** $i - \text{prev} \leq 2$ **then** ▷ index difference should not exceed 2 to be considered as consecutive
      $\text{APPEND}(H, h)$
    **else**
      $\mathcal{H}_s \leftarrow \mathcal{H}_s \cup \{\text{TUPLE}(H)\}$
      $H \leftarrow [\,h\,]$
    **end if**
    prev $\leftarrow i$
  **end for**
  $\mathcal{H}_s \leftarrow \mathcal{H}_s \cup \{\text{TUPLE}(H)\}$
  **return** $\mathcal{H}_s$
**end function**

**Algorithm S5** Find the most common tuple, weighted by length

**Require:** Multiset of tuples $\mathcal{H}$ ▷ duplicates are allowed
**Ensure:** Tuple $H_{\text{rep}}$

**function** MostCommon($\mathcal{H}$)
  $c \leftarrow$ empty map ▷ count unique tuples in $\mathcal{H}$
  $c_{\text{can}} \leftarrow$ empty map ▷ count unique canonical tuples in $\mathcal{H}$
  $\mathcal{H}_{\text{can}} \leftarrow \varnothing$ ▷ set of canonical tuples
  **for all** $H$ in $\mathcal{H}$ **do**
    **if** $H \notin c$ **then** $c(H) \leftarrow 0$
    $c(H) \leftarrow c(H) + 1$
    $H_{\text{can}} \leftarrow \min(H, \text{Reverse}(H))$ ▷ canonical tuple (lexicographical minimum)
    $\mathcal{H}_{\text{can}} \leftarrow \mathcal{H}_{\text{can}} \cup \{H_{\text{can}}\}$
    **if** $H_{\text{can}} \notin c_{\text{can}}$ **then** $c_{\text{can}}(H_{\text{can}}) \leftarrow 0$
    $c_{\text{can}}(H_{\text{can}}) \leftarrow c_{\text{can}}(H_{\text{can}}) + 1$
  **end for**
  $H_{\text{rep}} \leftarrow \arg\max_{H \in \mathcal{H}_{\text{can}}} |H| \cdot c_{\text{can}}(H)$ ▷ the most common canonical ordering, weighted by length
  $H'_{\text{rep}} \leftarrow \text{Reverse}(H_{\text{rep}})$
  **if** $c(H'_{\text{rep}}) > c(H_{\text{rep}})$ **then** $H_{\text{rep}} \leftarrow H'_{\text{rep}}$ ▷ choose orientation with greater support
  **return** $H_{\text{rep}}$
**end function**

#### Supplementary Note 1: Algorithmic Overview of Seqwin

Similar to recent $k$-mer-based approaches (Table 1), Seqwin allows the inclusion of “imperfect” $k$-mers: $k$-mers that are absent in some target genomes and/or present in some non-target genomes, making it robust to variations and errors in large datasets. However, Seqwin applies a novel minimizer graph algorithm that operates on a ∼1% sketch of all input $k$-mers, retaining linear time and space complexity and thereby scaling to tens of thousands of microbial genomes on modest hardware.

Seqwin first builds a weighted pan-genome minimizer graph, similar to the minimizer graph described by Coombe et al. (2020) but without the restriction that each minimizer should be present in all input genomes (Figure 1b). Each graph node represents a distinct minimizer observed in any genome, and each undirected edge connects two minimizers if they are found adjacent in at least one genome, weighted by the number of different genomes in which that minimizer adjacency occurs.

Second, Seqwin evaluates each node in the minimizer graph by computing a “penalty” score based on the L2 norm of its absence in target genomes and presence in non-target genomes. For example, a minimizer that is present in all target genomes and not present in any non-target genomes would have a penalty of 0, and a minimizer that is absent from all targets and present in all non-targets would have a penalty of $\sqrt{2}$. Thus, a series of consecutive low-penalty minimizers represents a genomic region that is both prevalent in target genomes and absent / dissimilar in non-target genomes. An example of a minimizer graph with node penalties is shown in Figure 1c.

Third, low-penalty subgraphs are extracted via seeded greedy breadth-first search (BFS) expansion (Figure 1c). Each subgraph consists of a set of connected nodes whose average penalty does not exceed a penalty threshold ($\tau_v$). This tolerates a small number of (relatively) high-penalty nodes in a low-penalty subgraph, resulting in larger subgraphs (longer signatures) and increasing search flexibility. The penalty threshold $\tau_v$ is arguably the most important parameter of Seqwin, distinguishing Seqwin from other tools with strict $k$-mer presence / absence criteria. However, $\tau_v$ should be set according to the homogeneity of input genomes. For example, for species with higher intra-species genomic homogeneity (e.g., *Mycobacterium tuberculosis*), a lower penalty threshold is preferred, while for other species such as *Salmonella enterica*, a higher penalty threshold might be preferred. In order to determine $\tau_v$ for arbitrary sets of input target and non-target genomes, we derive an intuitive method to calculate $\tau_v$ based on expected $k$-mer presence / absence, and estimate the expectations with Jaccard indices.

Lastly, Seqwin determines a representative minimizer ordering for each low-penalty subgraph, based on the most common minimizer ordering in target genomes (Figure 1d). This involves 1) finding the maximal consecutive occurrence of the subgraph's minimizers in each target genome, and 2) choosing the most common one across all target genomes, weighted by length. The genomic sequence corresponding to this minimizer ordering is the representative sequence, and is reported as the signature sequence. A more detailed illustration can be found in Figure S1. Alternatively, one could make a partial order alignment (POA) of the maximal occurrences from all target genomes, and determine the representative based on the consensus minimizer ordering of the POA.

Each candidate signature is evaluated for its sensitivity and specificity. Since both metrics are used to evaluate the performance of an assay (e.g., qPCR and dPCR), they can only be tested given an assay design (e.g., primers and probes) and a wet-lab setting. To estimate them in silico, Seqwin runs BLAST on each signature sequence against all target and non-target genomes, and summarizes BLAST results as two metrics: conservation and divergence, representing sensitivity and specificity, respectively. Conservation measures how consistently the signature sequence is conserved among target genomes, while divergence measures how dissimilar it is in non-target genomes (Supplementary Note 2). Seqwin outputs signature sequences with both high conservation and divergence as top candidates. Importantly, divergence is calculated based on the mismatches and gaps in the BLAST alignments, provided there are alignments in the non-target genomes. That is, those with no alignment (e.g., completely absent in non-target genomes) are not preferred, since they are more likely to be mobile genetic elements (MGEs), which have been shown to be highly problematic as signature sequences, given their ability to cut-and-paste or copy-and-paste into other bacterial genomes.

#### Supplementary Note 2: Evaluation of candidate signatures

Each output signature is aligned to all input genomes using NCBI BLAST+ (version 2.16.0). For each signature, its best BLAST alignment (highest `bitscore`) in each input genome is kept. The BLAST command and its arguments are shown below:

```
blastn -task blastn -max_hsps 1000 -max_target_seqs 50000
```

with `max_hsps` and `max_target_seqs` set to large numbers to keep all BLAST alignments.

Based on the BLAST alignments, we define two metrics to quantify sensitivity and specificity: conservation and divergence. Conservation measures how consistently the signature is conserved among target genomes, while divergence measures how dissimilar it is in non-target genomes. Formally, let the signature length be $L$. For all target genomes, we sum the number of identical bases (`nident` in BLAST outputs) in each BLAST alignment; let this sum be $I_{\text{target}}$. We define conservation as the average identity fraction in targets:

$$\text{conservation} = \frac{I_{\text{target}}}{L \cdot N_{\text{target}}} \tag{S1}$$

If a target genome has no BLAST alignment for the signature, it contributes 0 identical bases (thus lowering conservation). Thus, conservation = 1 would mean the signature is identical in all target genomes, whereas a lower value indicates some targets have mismatches in the alignment (or have no alignment).

Similarly, for all non-target genomes, we sum the number of nucleotide differences (`mismatch` and `gaps` in BLAST outputs) in each BLAST alignment. Let this sum be $D_{\text{non-target}}$. We define divergence as the average fraction of differences in non-targets:

$$\text{divergence} = \frac{D_{\text{non-target}}}{L \cdot N_{\text{non-target}}} \tag{S2}$$

It is important to note that the sum $D_{\text{non-target}}$ is only accumulated when there is a BLAST alignment. For a non-target genome with no BLAST alignment to the signature, it would make zero contribution to the sum. If the BLAST alignment is partial (e.g., only the first half of the signature is aligned), the flanking regions also have zero contribution to the sum. This way, a high divergence score indicates that the signature, while present in non-targets, has many differences, rather than being completely absent. This reduces the chance of picking up MGEs that might be completely absent in some non-target genomes. For each signature, the fraction of non-target genomes with a BLAST alignment can be found in Figures S3 to S5.

Finally, we compute a total score for each signature as the sum of its conservation and divergence. Seqwin uses this score to sort the output signatures so that those appearing first are the top candidates.

#### Supplementary Note 3: Calculating expected $k$-mer presence with Jaccard indices

Suppose we have two groups of genomes, and each genome is represented by a set of unique $k$-mers. Group $\mathcal{A}$ has $N$ $k$-mer sets and group $\mathcal{B}$ has $M$ $k$-mer sets. Note that $\mathcal{A}$ and $\mathcal{B}$ could be the same genome group.

$$\mathcal{A} = \{A_1, \ldots, A_N\},\ \mathcal{B} = \{B_1, \ldots, B_M\}$$

Suppose $A_i$ ($1 \leq i \leq N$) is a random genome in $\mathcal{A}$, and $h$ is a random $k$-mer in $A_i$. Here we want to calculate the expected number of genomes in $\mathcal{B}$ that have $h$. The Jaccard index between $A_i$ and $B_j$ ($1 \leq j \leq M$) is

$$J_{ij} = \frac{|A_i \cap B_j|}{|A_i \cup B_j|} = \frac{|A_i \cap B_j|}{|A_i| + |B_j| - |A_i \cap B_j|} \tag{S3}$$

Suppose all $k$-mer sets have the same size $s$, then

$$|A_i \cap B_j| = \frac{2J_{ij}}{1 + J_{ij}} \cdot s \tag{S4}$$

The probability that a random $k$-mer $h$ in $A_i$ is also found in $B_j$ is

$$\Pr(h \in B_j \mid h \in A_i) = \frac{|A_i \cap B_j|}{s} = \frac{2J_{ij}}{1 + J_{ij}} \tag{S5}$$

The expected number of $k$-mer sets in $\mathcal{B}$ that contain $h$ is

$$\mathbb{E}[\#\{B_j : h \in B_j\} \mid h \in A_i] = \sum_{j=1}^{M} \frac{2J_{ij}}{1 + J_{ij}} \tag{S6}$$

If $A_i$ is drawn uniformly from $\mathcal{A}$, then

$$\mathbb{E}[\#\{B_j : h \in B_j\}] = \frac{1}{N} \sum_{i=1}^{N} \sum_{j=1}^{M} \frac{2J_{ij}}{1 + J_{ij}} \tag{S7}$$

When $\mathcal{A}$ and $\mathcal{B}$ are identical and both represent the target genomes, we have

$$\begin{aligned} \mathbb{E}[\mathrm{f_t}(h)] = \frac{\mathbb{E}[\mathrm{F_t}(h)]}{|\mathcal{B}|} &= \frac{\mathbb{E}[\#\{B_j : h \in B_j\}]}{M} \\ &= \frac{1}{N^2} \sum_{i=1}^{N} \sum_{j=1}^{N} \frac{2J_{ij}}{1 + J_{ij}} \quad , \ (M = N) \end{aligned} \tag{S8}$$

When $\mathcal{A}$ is the set of target genomes and $\mathcal{B}$ is the set of non-target genomes, we have

$$\begin{aligned} \mathbb{E}[\mathrm{f_n}(h)] = \frac{\mathbb{E}[\mathrm{F_n}(h)]}{|\mathcal{B}|} &= \frac{\mathbb{E}[\#\{B_j : h \in B_j\}]}{M} \\ &= \frac{1}{N \cdot M} \sum_{i=1}^{N} \sum_{j=1}^{M} \frac{2J_{ij}}{1 + J_{ij}} \end{aligned} \tag{S9}$$

Definitions of $\mathrm{f_t}(h)$ and $\mathrm{f_n}(h)$ can be found in Equation (2) in the main text, which is also pasted below.

> ...suppose a minimizer node $h$ is found in $\mathrm{F_t}(h)$ target genomes and $\mathrm{F_n}(h)$ non-target genomes. Let
>
> $$\mathrm{f_t}(h) = \frac{\mathrm{F_t}(h)}{N_{\text{target}}}, \ \mathrm{f_n}(h) = \frac{\mathrm{F_n}(h)}{N_{\text{non-target}}}$$

Finally the penalty threshold can be calculated with Equation (4) in the main text, which is also pasted below.

> Consider a random $k$-mer $h$ (not necessarily a minimizer) sampled from a random target genome (select the target genome first and then select the $k$-mer). $1 - \mathrm{f_t}(h)$ is the fraction of target genomes that do not include $h$ (absence), and $\mathrm{f_n}(h)$ is the fraction of non-target genomes that include $h$ (presence), as defined in Equation (2). $\tau_v$ is then calculated with the expectations of these fractions
>
> $$\tau_v = \alpha_v \cdot \sqrt{(1 - \mathbb{E}[\mathrm{f_t}(h)]) \cdot \mathbb{E}[\mathrm{f_n}(h)]}$$
>
> which is the geometric mean of expected $k$-mer absence and presence, multiplied by a constant $\alpha_v$.

#### Supplementary Note 4: Filtering of the minimizer graph

To simplify the graph and remove artifacts caused by assembly errors or low-quality regions in the input genomes, low-weight edges are pruned based on a dynamic edge weight threshold. The purpose of this step is to reduce the graph and accelerate the breadth-first search (BFS) process. Therefore, only edges with negligibly low weights are removed and the original graph is preserved as much as possible.

Here we explain the calculation of the edge weight threshold. Since Seqwin focuses on minimizer nodes with penalties below $\tau_v$, from Equations (2) and (3) we have

$$\begin{aligned} \mathrm{F}(h) = \mathrm{F_t}(h) + \mathrm{F_n}(h) &\geq (1 - \mathrm{p}(h)) \cdot N_{\text{target}} \\ &\geq (1 - \tau_v) \cdot N_{\text{target}}, \text{ if } \mathrm{p}(h) \leq \tau_v \end{aligned} \tag{S10}$$

where $h$ is a minimizer node whose penalty is smaller than $\tau_v$, $\mathrm{F}(h)$ is the number of input genomes containing $h$. Since the weight of any edge incident to $h$ cannot exceed $\mathrm{F}(h)$ (an edge's weight is limited by the least frequent of its two nodes), the edge weight threshold is defined as

$$\tau_e = \alpha_e \cdot (1 - \tau_v) \cdot N_{\text{target}} \tag{S11}$$

where $\alpha_e$ is a small constant (0.3 by default). We prune any edges with weight less than $\tau_e$. The value of $\alpha_e$ is arbitrary and should not significantly affect the outputs, as long as it keeps $\tau_e$ significantly smaller than $(1 - \tau_v) \cdot N_{\text{target}}$, so that edges incident to low-penalty nodes are not pruned. After pruning low-weight edges, isolated nodes with degree of zero are also removed from the graph.

#### Supplementary Note 5: Benchmark settings

Benchmarking was performed on a Linux server (Red Hat Enterprise Linux release 8.9 (Ootpa)) with an AMD EPYC 7742 64-core processor (3.40 GHz) and 1 TB of memory.

Fur (`https://github.com/EvolBioInf/fur`, version 4.3, commit `168a9b6...c46ff95`) was compiled and installed from source. Dependencies of Fur were also installed: NCBI BLAST+ (version 2.16.0) and libdivsufsort (version 2.0.2) were installed via Bioconda; Phylonium (`https://github.com/EvolBioInf/phylonium`, version 1.7, commit `cf29672...36a57ad`) was compiled and installed from source. Fur was run under default parameters, using a single CPU thread (the *E. coli* dataset) or 20 CPU threads (the NCBI dataset). Specifically, a Fur database was first built for each target taxon, for example, for strain A in the *E. coli* dataset

```
makeFurDb -t only-A -n exclude-A -d A.fur -T 1
```

where `only-A` and `exclude-A` are directories containing FASTA files for target and non-target genomes, respectively, `A.fur` is the Fur database, and `-T` specifies the number of CPU thread(s). Fur would then search the database for signatures:

```
fur -d A.fur -t 1 > signatures-A.fasta
```

where `signatures-A.fasta` includes output signatures. Running time of Fur was calculated as the total time of both steps.

Unikseq (version 2.0.1) was installed via Bioconda, with $k$-mer length set to 21 (`-k 21`, same as Seqwin) and other parameters kept as default. For each target taxon, one of the genomes was chosen as the reference for Unikseq (noted in Supplementary Data 1-4). Genomes in the target and non-target groups were each merged into separate FASTA files. For example,

```
unikseq.pl -r ref.fasta -i tar.fasta -o neg.fasta -k 21
```

where `ref.fasta` is the reference, `tar.fasta` and `neg.fasta` are merged FASTA files of target and non-target genomes, respectively. Note that Unikseq did not support multithreading.

Neptune (version 2.0.0) was installed via Bioconda, with $k$-mer length set to 21 (`-k 21`, same as Seqwin) and other parameters kept as default. For each target taxon, one of the genomes was chosen as the reference (same as Unikseq, noted in Supplementary Data 1-4). For example,

```
neptune -i tar -e neg -o out -k 21 -r ref.fasta -p 20
```

where `tar` and `neg` are directories containing FASTA files for target and non-target genomes, respectively, `out` is the output directory, `ref.fasta` is the reference, and `-p` specifies the number of CPU process(es).

Seqwin (version 0.2.3, archived at https://doi.org/10.5281/zenodo.19056444) was run with the `--no-blast` flag to skip signature evaluation with BLAST, using a single CPU thread (the *E. coli* dataset) or 20 CPU threads (the NCBI dataset). For example,

```
seqwin --tar-paths tar.txt --neg-paths neg.txt --no-blast --threads 20
```

where `tar.txt` and `neg.txt` are lists of file paths to target and non-target genomes, respectively, and `--threads` specifies the number of CPU thread(s). For the full NCBI datasets *C. difficile* (3,995 genomes), *M. tuberculosis* (8,296 genomes) and *S. enterica* (14,822 genomes), the `--no-mash` flag was also added to use minimizer sketches to estimate penalty thresholds, instead of the default MinHash sketches (Mash).

#### Supplementary Note 6: Annotation of signatures

Reference genomes of the three target pathogens were downloaded from NCBI: *C. difficile* (GenBank accession `GCA_018885085.1`), *M. tuberculosis* (GenBank accession `GCA_000195955.2`) and *Salmonella enterica* subspecies *enterica* (GenBank accession `GCA_000006945.2`).

We first located the genomic location of each signature in its corresponding reference genome, using BLAST. All signatures had at least one BLAST hit in their corresponding reference genome. Since the reference genomes came with gene annotations, we extracted the gene overlapping with each signature. If more than one gene overlapped a signature, the one with the longest overlapping region was chosen. For each protein-coding gene, we extracted its amino acid sequence and submitted it to eggNOG-mapper (version 2.1.13) to get a more detailed annotation of its function, including its COG category, GO term, etc.

We also submitted the three reference genomes to the Mobilome Annotation Pipeline (MAP). MAP was one of the many pipelines under MGnify, and it was downloaded from its GitHub repository (https://github.com/EBI-Metagenomics/mobilome-annotation-pipeline, version 4.0.0, commit `9d9ec78...23f2653`). For each reference genome, MAP output a list of predicted mobile genetic elements (MGEs). We labeled signatures overlapping with any of the predicted MGEs.
